# Comparison of commercial RT-PCR diagnostic kits for COVID-19

**DOI:** 10.1101/2020.04.22.056747

**Authors:** Puck B. van Kasteren, Bas van der Veer, Sharon van den Brink, Lisa Wijsman, Jørgen de Jonge, Annemarie van den Brandt, Richard Molenkamp, Chantal B.E.M. Reusken, Adam Meijer

## Abstract

The final months of 2019 witnessed the emergence of a novel coronavirus in the human population. Severe acute respiratory syndrome coronavirus 2 (SARS-CoV-2) has since spread across the globe and is posing a major burden on society. Measures taken to reduce its spread critically depend on timely and accurate identification of virus-infected individuals by the most sensitive and specific method available, i.e. real-time reverse transcriptase PCR (RT-PCR). Many commercial kits have recently become available, but their performance has not yet been independently assessed.

The aim of this study was to compare basic analytical and clinical performance of selected RT-PCR kits from seven different manufacturers (Altona Diagnostics, BGI, CerTest Biotec, KH Medical, PrimerDesign, R-Biopharm AG, and Seegene).

We used serial dilutions of viral RNA to establish PCR efficiency and estimate the 95% limit of detection (LOD95%). Furthermore, we ran a panel of SARS-CoV-2-positive clinical samples (n=16) for a preliminary evaluation of clinical sensitivity. Finally, we used clinical samples positive for non-coronavirus respiratory viral infections (n=6) and a panel of RNA from related human coronaviruses to evaluate assay specificity.

PCR efficiency was ≥96% for all assays and the estimated LOD95% varied within a 6-fold range. Using clinical samples, we observed some variations in detection rate between kits. Importantly, none of the assays showed cross-reactivity with other respiratory (corona)viruses, except as expected for the SARS-CoV-1 E-gene.

We conclude that all RT-PCR kits assessed in this study may be used for routine diagnostics of COVID-19 in patients by experienced molecular diagnostic laboratories.

## INTRODUCTION

Coronavirus disease 2019 (COVID-19) is caused by the severe acute respiratory syndrome coronavirus 2 (SARS-CoV-2). This virus emerged in the human population in the final months of 2019 from a, so far unidentified, animal reservoir and has since spread across the globe (1). The SARS-CoV-2 pandemic poses an enormous burden on society, economic and healthcare systems worldwide, and various measures are being taken to control its spread. Many of these measures critically depend on the timely and accurate diagnosis of virus-infected individuals. Real-time reverse transcription polymerase chain reaction (RT-PCR) is the most sensitive and specific assay and therefore preferred (2, 3). Whereas many COVID-19 RT-PCR kits are currently commercially available, an independent assessment of these products is not yet publicly available and direly needed to guide implementation of accurate tests in a diagnostic market that is flooded with new tests. As of 11 April 2020, the FIND organization listed 201 molecular assays on their website as being on the market (www.finddx.org/covid-19/pipeline).

Coronaviruses are positive-stranded RNA viruses that express their replication and transcription complex, including their RNA-dependent RNA polymerase (RdRp), from a single, large open reading frame referred to as ORF1ab (4). The coronavirus structural proteins, including the envelope (E), nucleocapsid (N), and spike (S) proteins, are expressed via the production of subgenomic messenger RNAs, which during certain stages of the replication cycle far outnumber (anti)genomic RNAs. The ORF1ab/RdRp, E, N, and S genes are the targets most frequently used for SARS-CoV-2 detection by RT-PCR. For example, the “Corman” PCR, which was co-developed in our lab and is now routinely used for our in-house diagnostic work, targets a combination of the E-gene and the RdRp-gene (2). In this set-up, the E-gene primer/probe set is specific for bat(-related) betacoronaviruses, and therefore detects both SARS-CoV-1 and -2. In addition, whereas the RdRp-gene primers are also specific for bat(-related) betacoronaviruses, two probes are used: one specific for bat(-related) betacoronaviruses and another specific for SARS-CoV-2. In this study we only used the RdRp probe that is specific for SARS-CoV-2.

Here, we provide a comparison of a selection of seven readily available COVID-19 RT-PCR kits from different manufacturers (Table 1). One of these kits (BGI) was recently also included in a comparative study of various SARS-CoV-2 primer/probe sets (5). Most of the selected kits are CE-IVD certified and can be produced in large quantities. Using a dilution series of SARS-CoV-2 RNA we determine the 95% limit of detection (LOD95%) for each of these assays. In addition, a concise panel of clinical samples (n=22) was run to provide a first indication of clinical sensitivity and specificity. Although some kits appeared to perform better than others at identifying clinical samples at very low concentrations of SARS-CoV-2 RNA, all tests were able to identify positive samples with Ct≤34.5 in our in-house E-gene PCR. Therefore, we conclude that all of the RT-PCR kits assessed in this study may be used for routine diagnostics of COVID-19 by experienced molecular diagnostic laboratories.

**Table 1.** Overview of kits for RT-PCR-based detection of SARS-COV-2 included in the study.

| Manufacturer | Country | Catalog number | Storage condition | Regulatory status | Target gene(s) |
| --- | --- | --- | --- | --- | --- |
| Altona Diagnostics | Germany | 821003 | -20°C | RUO<sup>2</sup> | E<sup>1</sup>, S |
| BGI | China | MFG030010 | -20°C | CE-IVD | ORF1ab |
| CerTest Biotec | Spain | VS-NCO213L | RT | CE-IVD | ORF1ab, N |
| KH Medical | Korea | RV008 | -20°C | CE-IVD | RdRp, S |
| PrimerDesign | England | Z-Path-COVID-19-CE | -20°C<sup>3</sup> | CE-IVD | RdRp |
| R-Biopharm AG | Germany | PG6815RUO | -20°C | RUO<sup>4</sup> | E |
| Seegene | Korea | RP10244Y | -20°C | CE-IVD | RdRp, N, E<sup>1</sup> |

<sup>1</sup>As does the in-house "Corman" E-gene PCR, these E-gene assays are specific for both SARS-CoV-1 and -2.
<sup>2</sup>According to manufacturer's website the kit is RUO, the FindDx website states CE-IVD certification for this kit.
<sup>3</sup>Shipment is performed at RT
<sup>4</sup>According to the manufacturer, CE-IVD certification will be applied for in the near future.
Abbreviations: CE-IVD, European conformity label-in vitro diagnostics; E, envelope protein of SARS-CoV-2; RdRp, RNA-dependent RNA polymerase of SARS-CoV-2; N, nucleocapsid protein of SARS-CoV-2; ORF1ab, open reading frame 1a and b of SARS-CoV-2, includes the RdRp; RNA-dependent RNA polymerase of SARS-CoV-2, part of ORF1ab; RT, room temperature; RT-PCR, reverse-transcription polymerase chain reaction; RUO, research use only; S, spike protein of SARS-CoV-2; SARS-CoV-2, severe acute respiratory syndrome coronavirus 2.

## METHODS

### Selection of kits

Commercially available COVID-19 RT-PCR kits were identified via the FindDx website (www.finddx.org/covid-19/pipeline, March 2020) and requests for information and sample kits were sent via e-mail to approximately 20 manufacturers and/or distributors, focusing on those kits that had already obtained CE-IVD certification. Promising commercial kits were selected based on: 1) listing on the FindDx website; 2) responsiveness to requests; 3) accessible information (in English); 4) compatibility with different PCR platforms; 5) considerable production capacity. Notably, all of the PCR kits that we had selected for our analysis have in the meantime also been selected for the first round of independent evaluation by FIND (www.finddx.org/covid-19/sarscov2-eval-molecular/, April 2020). All of the kits included in our analysis were provided free of charge and none of the manufacturers were involved in the assessment and interpretation of the results. The selection encompasses both kits that require transport and storage at −20°C and kits that can be transported and stored at room temperature. Target genes for each RT-PCR kit were available in the assay documentation or upon request (for an overview, see Table 1). All PCRs were run on a LightCycler 480 II (LC480II, Roche) and performed according to the manufacturer’s instructions for use. Of note however, for some kits (BGI, KH Medical, and Seegene) settings for the LC480II were not provided and were therefore adapted from those provided for another machine.

### PCR efficiency and limit of detection

To establish PCR efficiency we first ran a duplicate 10-fold dilution series of viral RNA for each assay. Viral RNA was isolated from SARS-CoV-2 viral particles (hCoV-19/Netherlands/Diemen_1363454/2020, GISAID: EPI_ISL_413570) obtained from cell culture using the MagNA Pure LC Total Nucleic Acid Isolation Kit (Roche). We determined the slope by linear regression in GraphPad Prism and defined the required levels for PCR efficiency (E) and R^2^ as >95% and >0.95, respectively. Next, we ran four replicates of a 2-fold dilution series (diluted in yeast carrier RNA in water) to determine the LOD95% by Probit analysis using SPSS Statistics (IBM, version 24). The limited range of the dilution series did not allow for determination of a confidence interval for the LOD95% for all assays, which should therefore be regarded as an approximation and not considered definitive. The starting concentration of the viral RNA (copies/ml) was determined by digital PCR targeting the SARS-CoV-2 RdRp-gene and was specific for the positive sense genomic RNA (2).

### Clinical sensitivity and specificity

Finally, a panel of clinical samples with in-house confirmed SARS-CoV-2 (17.25≤Ct≤39.6 for the E-gene during initial diagnostics; n=16) or other respiratory viruses (influenza virus type A (n=2), rhinovirus (n=2), RSV-A and -B) was prepared (for Ct values obtained in initial diagnostics, see supplementary Table S1). RNA was isolated anew from stored clinical samples (naso- and/or oropharyngeal swabs in GLY-medium) using the MagNA Pure 96 DNA and Viral NA Small Volume Kit (Roche) and was assessed with a single replicate to obtain a first indication of clinical specificity and sensitivity. No re-test was performed when the result was inconclusive according to the manufacturer’s instructions for interpretation of the result (n=2). In addition to clinical samples, a panel of viral RNA from related cell cultured human coronaviruses (including SARS1, MERS, NL63, OC43, and 229E) was used to assess cross-reactivity within the coronavirus family (for Ct values of these samples see supplementary Table S1).

## RESULTS

### PCR efficiency was above the required level for all kits included in the study

We first assessed PCR efficiency for each target gene assay by running a duplicate 10-fold dilution series of SARS-CoV-2 viral RNA (Figure 1). All assays showed an efficiency ≥96% and R squares were >0.97, which are both well above the pre-defined required level. Since the applied filter settings were not correct for reading the Seegene N-gene assay, we excluded these data from all of our analyses.

**Figure 1.**
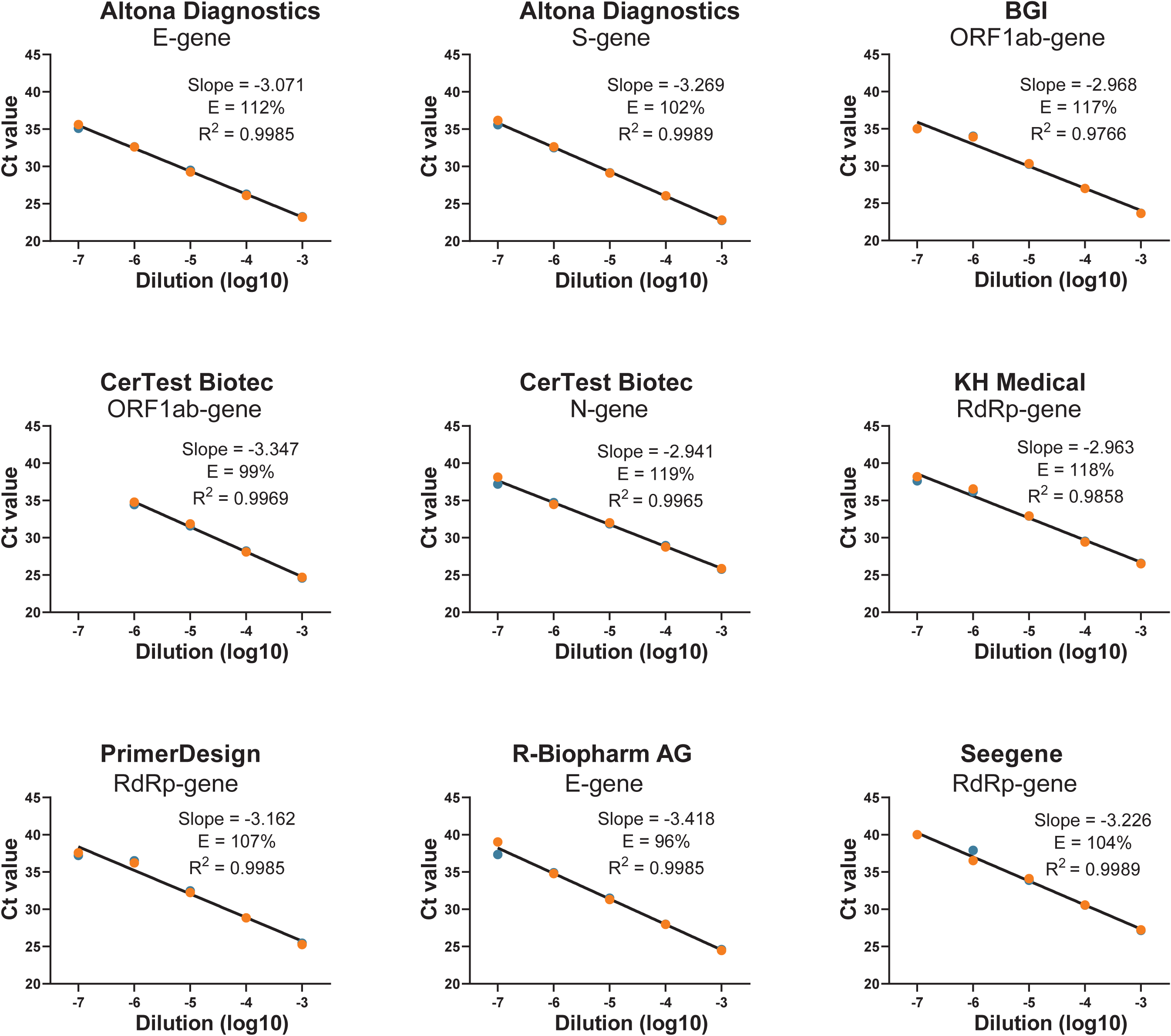
PCR efficiency for seven commercially available RT-PCR kits for the detection of SARS-CoV-2 RNA. PCR efficiency (E) for each target gene was assessed using a duplicate 10-fold dilution series of SARS-CoV-2 viral RNA. Linear regression was performed in Graphpad Prism to obtain the slope and R^2^. The percentage efficiency was calculated from the slope using the formula E = 100*(−1+10^−1/slope^). E-gene, gene encoding the envelope protein of SARS-CoV-2; RdRp, RNA-dependent RNA polymerase of SARS-CoV-2; N, nucleocapsid protein of SARS-CoV-2; ORF1ab, open reading frame 1a and b of SARS-CoV-2, includes the RdRp; RNA-dependent RNA polymerase of SARS-CoV-2, part of ORF1ab; S, spike protein of SARS-CoV-2; SARS-CoV-2, severe acute respiratory syndrome coronavirus 2.

### The LOD95% varied within a 6-fold range between the kits included in the study

The 10-fold dilution series provided a first indication of the LOD95% for each assay and were used to determine the starting point of a 2-fold dilution series performed with four replicates to come to a more precise estimate (for Ct values, see supplementary Table S2). Probit analysis was performed to estimate the LOD95%, which is shown in Table 2. Notably, due to the limited extent of the dilution series, this analysis did not always provide upper and lower bounds of the estimate and should not be considered definitive. We found that the estimated LOD95% for the various targets of the RT-PCR kits varied within a 6-fold range, with the RT-PCR kit from Altona Diagnostics having the lowest LOD95% at 3.8 copies/ml for both the E- and S-gene assays and the PrimerDesign kit having the highest LOD95% at 23 copies/ml (Table 2). Overall, our in-house “Corman” RT-PCR had the lowest estimated LOD95% at 0.91 copies/ml for the E-gene assay (2).

**Table 2.** Estimated limit of detection for SARS-COV-2 in copies/ml for individual assays.

| Company | LOD95% in copy/ml determined in this study<sup>1</sup> |  |  |  |
| --- | --- | --- | --- | --- |
|  | E | N | ORF1ab/RdRp | S |
| Altona Diagnostics | 3.8 (NA) | - | - | 3.8 (NA) |
| BGI | - | - | 4.3 (NA) | - |
| CerTest Biotec | - | 4.8 (NA) | 18 (13-56) | - |
| KH Medical | - | - | 4.8 (NA) | 4.3 (NA) |
| PrimerDesign | - | - | 23 (16-123) | - |
| R-Biopharm AG | 4.3 (NA) | - | - | - |
| SeeGene | 4.8 (NA) | NA<sup>2</sup> | 18 (13-56) | - |
| In-house PCR | 0.91 (0.61-2.4) | - | 3.1 (2.1-7.3) | - |

<sup>1</sup>The copy number was determined by digital PCR for the positive sense RdRp gene. Due to the limited range of the 2-fold dilution series, a confidence interval could not be determined for all assays.
<sup>2</sup>The filter settings for the Seegene N-gene PCR were not correct and these results are therefore excluded.
Abbreviations: E, envelope protein of SARS-CoV-2; LOD95%, 95% limit of detection; N, nucleocapsid protein of SARS-CoV-2; NA, not available; ORF, open reading frame; RdRp, RNA-dependent RNA polymerase of SARS-CoV-2; RT-PCR, reverse-transcription polymerase chain reaction; S, spike protein of SARS-CoV-2; SARS-CoV-2, severe acute respiratory syndrome coronavirus 2.

### The clinical sensitivity appears to vary between the kits included in the study

Next, we analyzed a panel of clinical samples previously submitted for routine SARS-CoV-2 diagnostics (n=16) for which the presence of various amounts of SARS-CoV-2 RNA had been confirmed using our in-house PCR. In addition, we included a panel of clinical samples (n=6) with other confirmed respiratory viral infections, including influenza virus type A, RSV A and B, and rhinovirus. Notably, the new RNA isolation performed on stored clinical samples resulted in increased Ct values (by approximately 1 Ct) compared to the initial diagnostic results for our in-house E-gene PCR. For this reason, even using our in-house PCR we could not confirm the presence of SARS-CoV-2 RNA in 3 out of 16 samples (see Figure 2A and supplementary Table S1). The positive identification rate for the various RT-PCR kits varied from 10 to 13 out of 16 samples (Figure 2A), with R-Biopharm AG performing best (13/16), followed by BGI, KH Medical, and Seegene (12/16), CerTest BioTec (11/16), and Altona Diagnostics and PrimerDesign (10/16). Of note, Seegene had one “inconclusive” sample according to the manufacturer’s instructions for interpretation, which might have tested positive upon re-testing but has now been counted as “negative”. All target gene assays were able to positively identify the 10 clinical samples with the highest concentrations of SARS-CoV-2 (Ct≤34.50 in our in-house E-gene PCR). For these samples, the different assays showed a similar pattern of Ct values, on average ranging from almost 1 Ct lower (Altona Diagnostics S-gene) to almost 5 Ct higher (KH Medical S-gene) than those obtained with the in-house E-gene PCR (Figure 2B).

**Figure 2.**
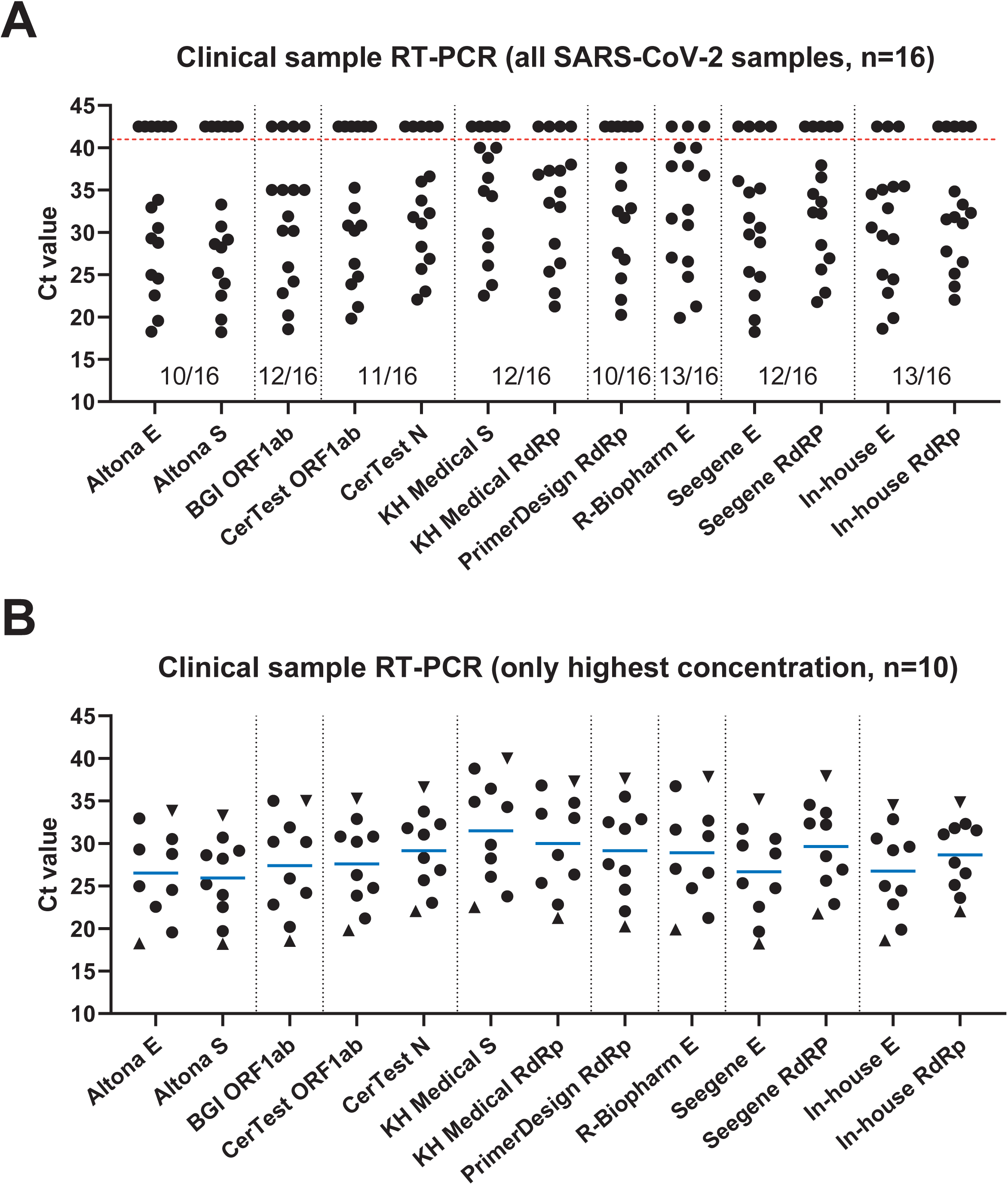
Different RT-PCR kits showed variations in detection rate and Ct values. RNA isolated from stored SARS-CoV-2-positive clinical samples using the MagNA Pure 96 DNA and Viral NA Small Volume Kit (Roche) was subjected to the various RT-PCR assays according to the manufacturer’s instructions for use, on a LightCycler 480 II (Roche). **A)** Graph depicts Ct values obtained for all clinical samples (n=16) in all RT-PCR assays. Data points above the red dotted line are negative, for plotting purposes indicated with Ct 42.5. The detection rate of the complete RT-PCR kit is indicated below the data points, e.g. 10/16 means 10 out of 16 samples tested positive according to the instructions for data interpretation provided by the manufacturer. For both the CerTest and Seegene kits, one sample was “inconclusive” according to the manufacturer’s guide for interpretation and was therefore counted as “negative”, although a signal was observed for at least one target. **B)** Graph depicts only data for those clinical samples (n=10) with the highest concentration of SARS-CoV-2 RNA and which were positively identified by all RT-PCR assays. The blue line shows the mean Ct value for each assay, triangles show the Ct values of the samples with the highest (sample 1) and lowest (sample 10) concentration according to the in-house E-gene PCR. E, envelope protein of SARS-CoV-2; RdRp, RNA-dependent RNA polymerase of SARS-CoV-2; N, nucleocapsid protein of SARS-CoV-2; ORF1ab, open reading frame 1a and b of SARS-CoV-2, includes the RdRp; RNA-dependent RNA polymerase of SARS-CoV-2, part of ORF1ab; S, spike protein of SARS-CoV-2; SARS-CoV-2, severe acute respiratory syndrome coronavirus 2.

### None of the assays showed cross-reactivity with circulating respiratory (corona)viruses

Importantly, none of the assays resulted in a positive signal for any of the clinical samples with confirmed non-coronavirus respiratory viral infections (Supplementary Table S1). We also ran a panel consisting of cell culture-derived viral RNA for related human coronaviruses (SARS1, MERS, NL63, OC43, and 229E) to check for cross-reactivity within the coronavirus family. Of these, only the SARS-CoV-1 E-gene was identified, as per design, by assays from Altona Diagnostics, Seegene, and our in-house PCR (Supplementary Table S1).

## DISCUSSION

Here we provide a comparison of seven commercially available RT-PCR kits for the detection of SARS-CoV-2 in clinical samples. All RT-PCR kits performed satisfactorily regarding PCR efficiency (≥96%) and the estimated LOD95% varied within a 6-fold range between kits (3.8-23 copies/ml). Notably, the copy number concentration of the standard was determined by digital PCR on the positive sense RdRp gene and therefore provides an indication of the number of viral particles per ml. The actual copy number for each RT-PCR target and accompanying limit of detection may vary depending on, for example, the amount of subgenomic messenger RNA-containing cells that are present in the (clinical) sample.

From a selection of clinical samples with various concentrations of viral RNA, all RT-PCR kits were able to positively identify the ten samples with the highest concentrations of SARS-CoV-2 RNA (Ct≤34.5 in our in-house E-gene PCR). To provide an indication on clinical relevance of this finding: from our in-house diagnostic data on patients presenting with COVID-19 symptoms, it appears that from all individuals testing positive for our in-house E-gene PCR (n=416) the proportion of individuals with a Ct value >34.5 is approximately 3.6% (unpublished data). The R-Biopharm AG kit positively identified the highest number of clinical samples, i.e. 13 out of 16, comparable with our in-house PCR. Three kits were able to positively identify 12 out of 16 samples (BGI, KH Medical, Seegene). Notably, we performed our analysis using only a small number of clinical samples and we therefore advise that diagnostic laboratories in the field conduct additional and more extensive in-house clinical validations upon implementation of novel RT-PCR kits. Importantly, none of the assays showed cross-reactivity towards a panel of other respiratory (corona)viruses, except for the expected cross-reactivity with the SARS-CoV-1 E-gene. Since the latter virus is no longer known to be circulating in the human population, we consider this cross-reactivity acceptable.

Considering our findings, we believe that all of the commercially available RT-PCR kits included in this study can be used for routine diagnostics of symptomatic COVID-19 patients. When performing virus diagnostics in populations that may be expected to display low viral loads, such as health-care workers with mild or no symptoms or patients during later stages of the infection (6), it might be advisable to use those kits that performed best regarding the positive identification of clinical samples, i.e. RT-PCR kits from R-Biopharm AG, BGI, KH Medical, and Seegene.

## ACKNOWLEDGEMENTS

We kindly thank Barbara Favié (RIVM, Bilthoven) for performing the digital PCR. In addition, we would like to thank all manufacturers who kindly donated their RT-PCR kits for our evaluation.

